# Downregulation of Keratin Proteins in the Neuroendocrine System of Amyotrophic Lateral Sclerosis with PT150 as a Dual Neuroprotectant

**DOI:** 10.1101/2025.03.18.643927

**Authors:** Samarth Dunakhe, Arnav Sharma, Aryav Das, Pranav Kulkarni

**Affiliations:** Department of Computational Neuroscience, CortexPD Labs; Troy, USA; School of Veterinary Medicine, University of California, Davis; Davis, USA

**Keywords:** Graph Neural Network, Amyotrophic Lateral Sclerosis, Diagnosis, Biomarkers, Keratin Proteins, Neuroendocrine System, PT150, Neuroprotectant

## Abstract

Amyotrophic Lateral Sclerosis (ALS) is a neurodegenerative disease characterized by motor neuron degeneration. Although ALS biomarkers exist, they lack specificity and early-onset. ALS also has a 40% misdiagnosis rate. This paper proposes the use of a Graph Neural Network (GNN) to diagnose ALS, new ALS-specific biomarkers, and a therapeutic based on this group. Gene expression and proteomic datasets were sourced. A kNN data representation was used to train a GNN for disease diagnosis. Achieving 98% accuracy between ALS and control, and 85% between ALS patients and ALS mimics (the specific diseases within this class were not disclosed), outperforming existing approaches. Integrated Gradients then quantified each genes importance in the model’s decision. KRTAP3-1 and KRTAP9-8 were substantial in the model’s classification process due to involvement in cytoskeletal organization. In parallel, statistical proteomi analysis revealed the significance of KRT6A, KRT6B, and KRT4 due to involvement in cytoskeletal organization, tissue development, and cellular differentiation, all related to ALS. These keratin proteins were expressed in the pituitary gland, suggesting that their downregulation contributes to hormonal dysregulation in ALS. Once Glucocorticoid Receptor (GR) was found to be the reason for keratin downregulation, molecular docking identified PT150 as a therapeutic, binding with and inhibiting GR and Cyclooxygenase 2, also reducing neuroinflammation in ALS.

## 1 Introduction

ALS is a neurodegenerative disease characterized by the progressive deterioration of upper and lower motor neurons that facilitate movement. Upper motor neurons (UMNs) are located in the motor cortex and brain stem along with lower motor neurons (LMNs) being in the spinal cord and brain stem. Degeneration of these motor neurons leads to functional impairments such as muscle atrophy, paralysis, and eventually respiratory failure.

ALS is one of the most heterogeneous neurodegenerative diseases due to its genetic variability, as only 15% of ALS cases are known to have genetic origin [11], ALS cases also have varying forms of onset such as bulbar and spinal which account for varying progression rates and varying mechanisms involved in ALS. This accounts for the 40% misdiagnosis rate and delayed treatment for ALS. Coupled with the 2-4 year average lifespan all ALS patients, this issue becomes critical to address [22].

Recent research has attempted to reduce this misdiagnosis rate through the discovery of several biomarkers.

- Neurofilament Light (NfL): A protein located in the cerebrospinal fluid classified as a biomarker when it is too abundant [21].
- TAR DNA-binding protein 43 (TDP-43) & RNA-binding protein FUS (FUS): A protein and encoding gene respectively, involved in DNA binding processes, contributing to ALS pathology when they are found in the incorrect cellular location (cytoplasm rather than nucleus) [18] [10].
- C9orf72: A protein involved in intracellular matrix transport [16].
- SOD1 Superoxide Dismutase (SOD1): An encoding gene classified as a biomarker when it encodes enzymes with toxic abilities [9].

These biomarkers have been validated to help researchers understand ALS pathology and improve diagnosis. However, these established biomarkers are nonspecific between other diseases along with ALS such as:

- NfL is observed in several cases of Alzheimer’s disease (AD), frontotemporal dementia (FTD), multiple sclerosis (MS), and Guillain-Barré syndrome (GBS).
- TDP-43 is observed in 90% of Hippocampal Sclerosis (HS) cases, 57% of Alzheimers Disease (AD) cases, 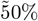 of Frontotemporal Dementia (FTD) cases [8] [7].
- FUS aggregates is observed in only 3-4% of familial ALS cases, only 1% of sporadic ALS cases, and 6-20% of Frontotemporal Lobar Degeneration (FTLD) cases [10] [5] [6].
- C9orf72 is observed in 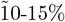 of FTD cases and 10% of ALS cases [15] [17].
- SOD1 is observed in only 10-20% of ALS cases [2] [3] [4].

Researchers have applied computational approaches to diagnose ALS, however, these approaches plateau at low accuracies around 80% [19] due to their limited data and suboptimal feature aggregation.

Observing these deficiencies, this paper surpasses the state-of-the-art method for diagnosing ALS using Graph Neural Networks, discovers new ALS-specific biomarkers, and a multi-target neuroprotective therapeutic.

## 2 Materials and Methods

### 2.1 Data Profile

The first data set was a batch-controlled whole-blood gene expression profile with 1117 subjects from Europe (397 ALS, 645 control, 75 mimics). The ALS patients had various forms disease onset, split between Bulbar or Spinal onset. Stabilized with PAXgene Tubes, 29,830 transcripts were expressed. [19].

The second dataset was a cerebrospinal fluid proteomic analysis with 101 subjects from the USA (35 sporadic ALS, 10 C9orf72 ALS, 6 SOD1 ALS, 6 asymptomatic C9orf72, 44 age-matched controls). Collected by tandem mass tag mass spectrometry (TMT-MS), the expression for 2,105 proteins was quantified [20].

The validation dataset was a gene expression profile for Alzheimer’s Disease (AD) with postmortem prefrontal cortex (PFC) samples from 467 subjects (310 AD and 157 non-demented control). The age in the dataset ranges from 53-100 years for the AD subects and 22–106 years for the control subjects. The dataset consists of 209 females, 258 males, and 19,206 transcripts [1].

The integration of transcriptomic and proteomic data allowed us to understand molecular perspectives across the central-peripheral disease axis. Using different patients (European whole-blood and USA cerebrospinal fluid) strengthened the analysis by providing external validation rather than introducing a single-dataset bias. Furthermore, blood gene expression is accessible for clinical monitoring, while CSF proteins capture neurodegeneration in the CNS. This approach was very strong as each dataset constantly validated eachother throughout the study.

### 2.2 Data Representation

The dataset was transposed so that each row corresponded to an individual patient. Diagnoses were encoded as 0, 1, or 2, representing control, ALS, and ALS mimic subjects, respectively. Then, an 80/20 train-test split was applied and all features were standardized using *StandardScaler*.

As the gene expression profile consisted of 29,830 features resulting in model overfitting, Principal Component Analysis simplified the dataset to 300 of its most considerable features using covariance calculations.

These 300 features were then used to construct the kNearest Neighbor graph where each node represented a patient sample (ALS, ALS mimic, control) and each edge connected each node to its 5 nearest neighbors, or 5 most similar samples. To preserve each nodes’ features during message passing, improve gradient flow during training, and to prevent over-smoothing, self loops were added. Self loops are connections that link nodes back to themselves, creating circular edges in the graph structure that regularize graph operations by adding their identity matrix to the adjacency matrix. This kNN was then mapped to a 256-dimensional space with a linear transformation. The resulting adjacency matrix was computed using kneighbors_graph() from scikit-learn, and the edges were formatted into a PyTorch Geometric tensor. These two parameters formed the input for the Graph Neural Network (GNN).

The GNN was trained using the Adam optimizer with a learning rate of 0.005 and weight decay of 5e-4 to prevent overfitting. The training process was set to take 50 epochs with equal class weights in the cross-entropy loss function, allowing the model to gradually learn the patterns in the data. The cross-entropy loss function is detailed as below:

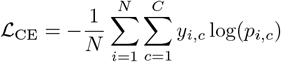

_ℒCE_ = Cross-entropy loss

*N* = Number of samples

*C* = Number of classes

*y*_*i,c*_ = Binary indication of accuracy

*p*_*i,c*_ = Predicted probability of classification

### 2.3 GNN Architecture

GraphSAGE (Graph Sample and Aggregate) is a GNN architecture designed for inductive learning. This allows the GNN to generate dynamic node embeddings with information from their neighbors in the feature space, described as “Neighborhood Aggregation”, rather than relying on fixed feature vectors. GraphSAGE works by sampling a fixed number of neighbors and applying functions (such as mean and pooling) to combine their features allowing each node to capture local and global patterns in the data when aggregated [13]. The GNN architecture for this task is described in Figure 1.

**Figure 1:**
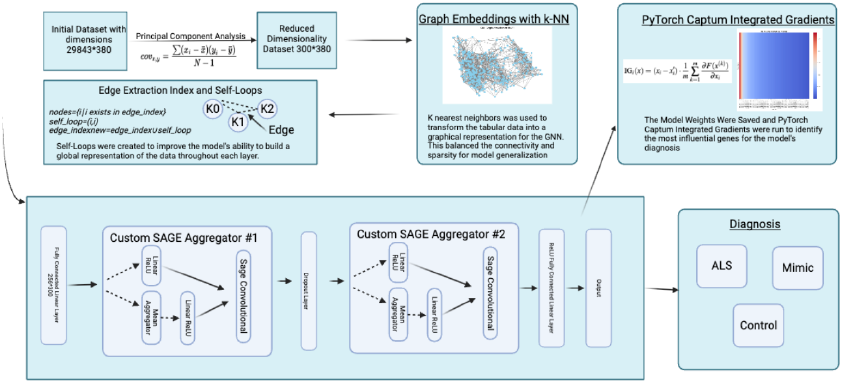
The Graph Neural Networks Architecture showing its complete processing pipeline.[27]

The **first SAGEConv layer** aggregated the data that was represented by the kNN graphs adjacency matrix and edges using neighborhood aggregation computed as:

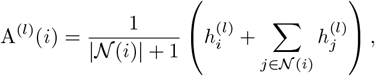

A^(*l*)^(*i*) = Aggregated feature representation

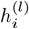= Feature representation of node *i* at layer *l*

*N* (*i*) = Set of neighboring nodes

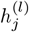= Representation of a node *j* at layer *l*

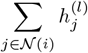= Summation of all node neighbors

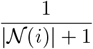= Normalization factor for mean aggregation

The **hidden layer** refined the node representations by applying Dropout regularization where 50% of the neurons were randomly set to zero druing training to prevent overfitting. It also employed ReLU activation enabling non-linearity for the graphs operations.

The **second SAGEConv layer** further aggregated each nodes information allowing the model to incorporate the secondary neighbors for each node in the feature space, described as “Heirarchical Aggregation” [13]. The resulting 256-dimensional node embeddings understood both the local and global graph structure.

The **fully-connected output layer** transformed the 256-dimensional node embeddings into classification outputs through a linear transformation into logits corresponding to each class. These logits were then converted to probability distributions using a softmax function mathematically represented below:

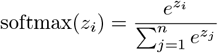

softmax(*z*_*i*_) = The softmax function applied to element *z*_*i*_.

*z*_*i*_ = An element of the input logits vector.

*n* = The number of elements in the vector

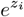= The exponential of the input element *z*_*i*_.

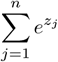= The sum of exponentials of the vectors elements

### 2.4 Feature Attribution

Integrated Gradients (IG) attributed each inputs relative importance in the model output.

IG works by interpolating between a baseline input *x*^′^ which represents the mean of the feature values across the data samples and the actual input. This generates a series of intermediate inputs that trace a line in the feature space. At each step, the gradient of the model’s output for each feature is computed, capturing how each input contributes to the models output [12]. The resulting equation, that approximates the integral with a summation, is given by:

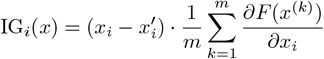

IG_*i*_(*x*) = Attribution for feature *i* at input *x*.

*x*_*i*_ = Actual value of feature *i*

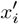= The mean of the feature values *i*,

*m* = Number of interpolation steps

*x*^(*k*)^ = Intermediate input

*F* (*x*) = Model’s prediction function

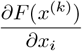= Gradient of the model’s output

Through this approach, we identified the genes most influential for the GNN when choosing to diagnose a patient with ALS.

### 2.5 Model Validation

To quantify the performance of the GNN, we used accuracy score, which measured the proportion of correctly classified samples to incorrectly classified samples and Area Under the Receiver Operating Characteristic curve (AUC-ROC) which evaluated the model’s confidence between classifications (ALS vs. control, ALS vs. mimics). Precision, recall, and F1 scores assessed the quality of the GNNs classification with consideration for the dataset’s class imbalance. Confusion matrices then visualized the True positive, true false, false positive, and false negatives in the classifications. The epoch and feature amount was changed to 100 and 250 respectively as it had the best performance as compared to 50 epochs and 300 features.

The GNN was cross-validated by assesing its accuracy on an Alzheimer’s gene expression profile to evaluate its generalizability. The pipeline was the same as it was on the ALS data with the only difference being how PCA reduced the dataset to 250 features rather than 300.

### 2.6 Statistical Analysis

Statistical analysis was conducted between the ALS and control samples in the proteomic data (n = 51 ALS samples, n = 44 control samples). We usedLog_2_-fold change (Log_2_FC) and two-sample t-tests to assess statistical significance and to quantify the relative protein abundance of each protein. Proteins were considered differentially expressed when they met both thresholds of (p < 0.05) and (|Log_2_FC| > 0.58).

### 2.7 Pathological Analysis

Analysis with UniProt and Gene Ontology (GO) was then performed to clarify which biological pathways and processes the 40 most differentially expressed proteins and genes were involved in [14] [24]. We identified major pathways, in common between the proteins and genes, with the p value cutoff at 0.05.

The 40 most differentially expressed proteins and genes were then neurospatially analyzed using the Allen Brain Atlas (ABA) and the Human Protein Atlas (HPA) [23] [25]. ABA and HPA provided the expression levels for each protein and gene across the brain along with their tissue-level specificities.

### 2.8 Therapeutic Discovery

Through this analysis, we identified a 3 biomarkers as our therapeutic targets. By observing existing literature, a specific receptor emerged as a binding target to normalize these 3 biomarkers. However, we need to find a molecule to bind with this receptor and normalize these 3 biomarkers.

First 10,000 molecular sequences were selected from PubChemPy in the form of SMILE sequences. Filtering was then conducted based on Lipinski’s rule of 5, which is a set of rules that determine whether a drug can be orally active:

- There are no more than 5 hydrogen bond donors
- There are no more than than 10 hydrogen bond acceptors
- The molecular mass is less than 500 daltons
- The octanol-water partition coefficient does not exceed 5

The resulting molecules were then put into the *Protein Ligand Binding Affinity Using Pretrained Transformers* (PLAPT) model, which uses pre-trained Transformers and a Branching Neural Network (BNN) to predict protein-molecule binding affinities with State-Of-The-Art accuracies [28].

The resulting molecule was put into Autodock Vina to validate its high binding affinity. [29]. Way2Drug was then used to assess Blood Brain Barrier Permeability, a major criteria for neurologically active drugs. [31]

## 3 Results

### 3.1 GNN Evaluation

The GNN performed at a 98% test accuracy + 99% AUC between ALS and control as shown in Figure 2a and a 85% test accuracy + 84% AUC between ALS and Mimics as shown in Figure 2b. This reveals the GNN’s ability to diagnose ALS with state-of-the-art accuracy.

**Figure 2:**
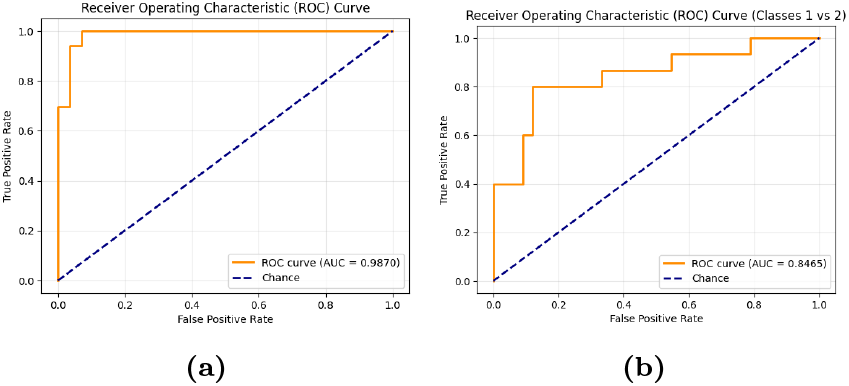
AUC-ROC graphs showing the trade-off between true positives and false positives for both classification tasks.

The confusion matrix on the left in Figure 3 (ALS vs Control Classification) shows that control subjects were classified with 28 true positives and 0 false negatives, while ALS subjects were classified with 28 true positives and 1 false negative (misclassified as Control). This shows near-perfect state-of-the-art classification performance for this task.

**Figure 3:**
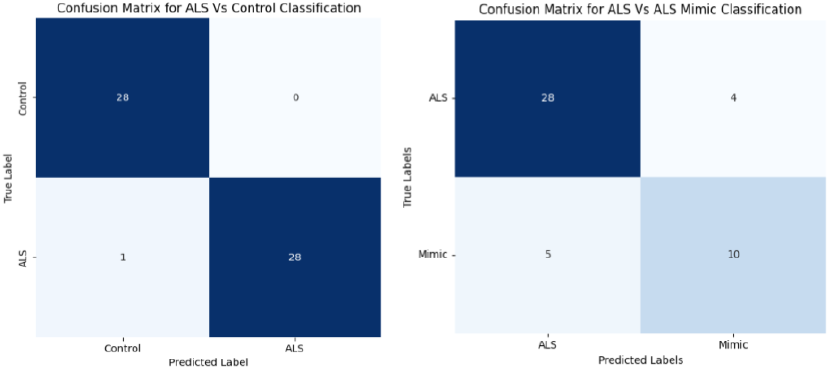
The confusion matrices showing the TP/TN/FP/FN rate for both classification tasks.

The confusion matrix on the right in Figure 3 (ALS vs ALS Mimic Classification) shows that ALS subjects were classified with 28 true positives and 4 false negatives (misclassified as Mimic), while Mimic subjects were classified with 10 true positives and 5 false positives (misclassified as ALS), showing high accuracies once again for this task. The GNN showed extreme generalizability with 97% test accuracy after 100 epochs. The GNN achieved balanced precision and recall values (0.94-1.00) in both classes, with an almost perfect AUC-ROC score of 0.98. The confusion matrix showed that only 2 out of 63 test samples were misclassified, confirming the GNNs ability to model complex relationships in the data and suggesting applications to diagnose several neurodegenerative diseases based on the data used to train the model.

### 3.2 Keratin Biomarkers

Integrated Gradients showed that the genes KRTAP3-1 and KRTAP9-8 were vital in the model’s decision making process, contributing to a relatively substantial portion of the model’s classification process. Similarly, in the proteins encoded by KRTAP3-1 and KRTAP9-1, downregulation was identified in the keratin proteins KRT6A, KRT6B, and KRT4 through the statistical test results shown in Table 2. Through pathological analysis with Gene Ontology and Uniprot, it was discovered that the keratin proteins and genes were involved in the pathways “tissue development”, “cytoskeletal organizaiton”, and “cellular differentiation”. These biomarkers display strong potential to improve the diagnostic accuracy of ALS and to provide insights into disease pathology.

**Table 1:** All models were trained on the same dataset. Existing machine/deep learning methods for ALS diagnosis on our dataset.

| Method | AUC ALS vs. Controls (%) | AUC ALS vs. Mimics (%) | Reference |
| --- | --- | --- | --- |
| Support Vector Machine (SVM) | 87 | 65 | [19] |
| Nearest Shrunken Centroids | 87 | 66 | [19] |
| LASSO | 90 | 68 | [19] |
| LASSO | 86.5 | — | [32] |
| Graph Neural Network (GNN) | **97.0** | 84 | Our work |

**Table 2:** Downregulation quantified of keratin protein variants in ALS patients (Log_2_FC < *−*0.8) with statistically significant *p*-values (*p <* 0.05).

| Variant | Log<sub>2</sub>FC | *p*-value |
| --- | --- | --- |
| KRT6B | −0.999 | 0.024 |
| KRT6A | −0.978 | 0.034 |
| KRT4 | −0.839 | 0.039 |

*Note:* Negative Log<sub>2</sub>FC values indicate downregulation.

Neurospatial analysis identified substantial keratin protein downregulation within the pituitary gland, a region implicated in the dysregulation of growth hormone and cortisol in amyotrophic lateral sclerosis (ALS) [33] [34]. As shown before, Uniprot analysis revealed that the 3 keratin proteins plays a critical role in cytoskeletal organization. The observed downregulation of keratin proteins in the pituitary gland suggests potential cellular instability within this region. These findings indicate that keratin proteins may contribute to hormonal dysregulation in ALS through cytoskeletal instability. This pathway is further shown in Figure 4.

**Figure 4:**
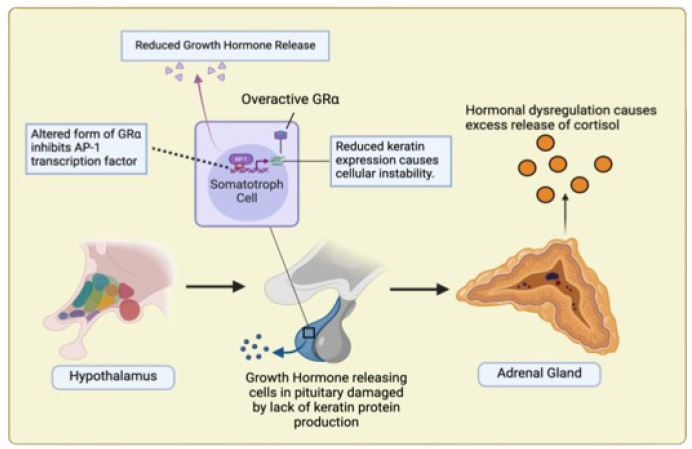
The pathway of the hormone dysregulation, specifically in IGF-1, in ALS. [30]

### 3.3 PT150 Neuroprotectant

To identify a therapeutic viable to regulate these keratin proteins and the subsequent hormonal dysregulation, we identified the 2 receptor targets of Glucocorticoid Receptor and Cyclooxygenase 2 as shown in Figure 5.

**Figure 5:**
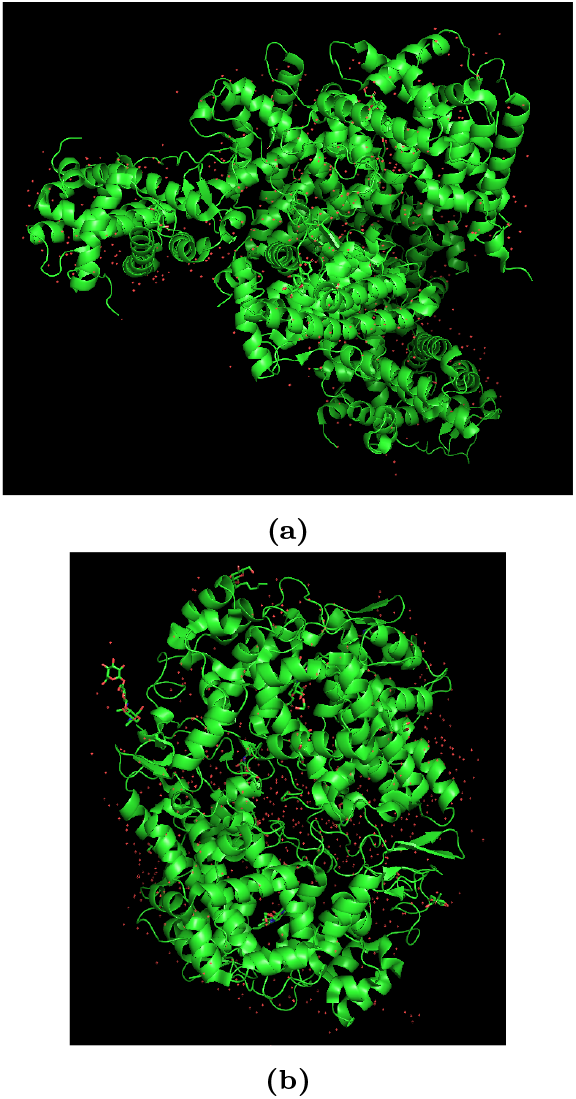
Crystal structures of cortisol-bound glucocorticoid receptor ligand binding domain (left) and aspirin-acetylated human cyclooxygenase-2 (right). [24]

- Glucocorticoid Receptor (GR) was shown to inhibit the activity of the AP-1 transcription factor, which is supposed to regulate keratin protein expression in response to stimuli such as neuronal activity and growth factors.
- Cyclooxygenase 2 (COX2) was identified as an important enzyme involved in the inflammatory response of ALS motor neuron degeneration. Specifically, COX2 produces prostaglandins, which cause inflammation and pain.

Through molecular docking with PLAPT, the molecule, PT150, was identified as a small-molecule modulator for GR activity, further substantiated by past literature detailing this interaction [35]. As shown in Figure 6, PT150 was able to bind with GR at -7.5 kcal/mol directly activating the AP-1 transcription factor. This, in turn, allowed for the AP-1 transcription factor to regulate KRT6A, KRT6B, and KRT4 regulating hormonal release in ALS, especially in IGF-1 production. Furthermore, PT150 was able to bind with Cyclooxygenase 2 (COX 2) at -7.5 kcal/mol which is the enzyme responsible for neuroinflammation. This is important in ALS as neuroinflammation directly contributes to motor neuron degeneration through its application of oxidative stress, release of cytokines, activation of microglia cells, and disruption of the Blood-Brain-Barrier (BBB).

**Figure 6:**
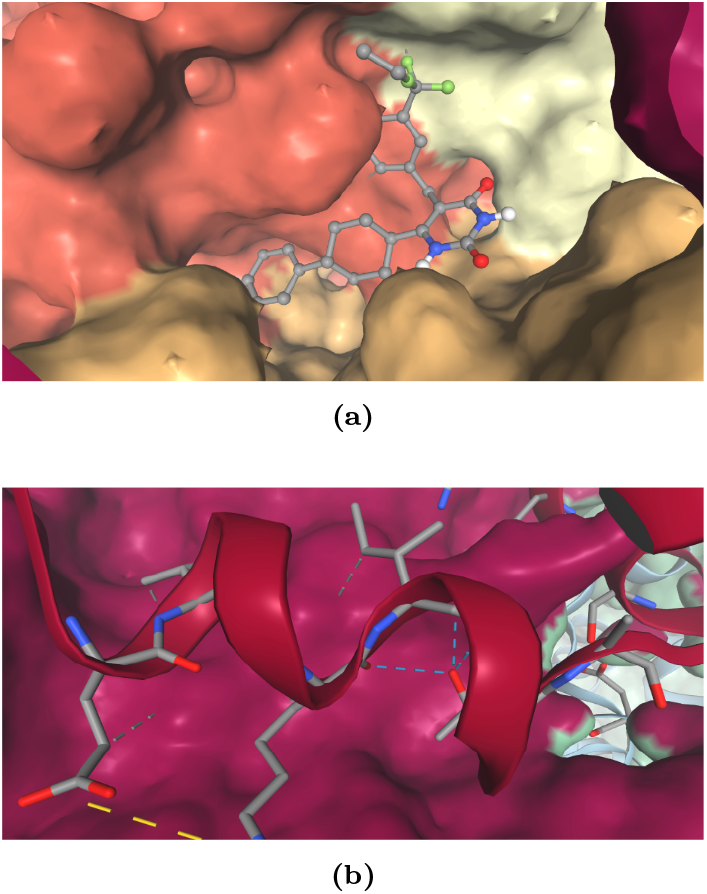
Simulated binding environments between PT150 to Glucocorticoid Receptor and Cyclooxygenase 2 both binding at -7.5kcal/mol. [29]

Way2Drug displayed PT150’s BBB permeability, as shown by its LogBB value of -0.04, indicating that it will be present in similar concentrations within the brain and the blood.

## 4 Discussion

From a **data standpoint**, longitudinal sampling with a more diverse population from more continents would help observe if KRT6A, KRT6B, and KRT4 could act as population-generalized ALS biomarkers.

From a **clinical standpoint**, validation of cellular-level roles of the keratin proteins in ALS can be carried out through genetic engineering (eg CRISPR Cas9), in vivo assays using tissue-level expression assays such as Western blotting, and behavioral analysis of KRT6A, KRT6B, and KRT4 dysregulation in animal models.

From a point of view of **therapeutic development**, PT150 has already been clinically applied as a therapeutic in its Phase II trials to treat depression, but its ability to regulate KRT6A, KRT6B and KRT4 can be investigated using mouse models with ALS SOD1 to assess hormone regulation and techniques such as immunofluorescence or qPCR complemented with Western Blotting to quantify and confirm regulation of the keratin protein.

The threat of **keratin contamination** was addressed in this research. The proteomic dataset stated “MS/MS spectra were searched against all canonical Human proteins downloaded from Uniprot (20,402; accessed 02/11/2019), as well as common contaminants (51 total)” [20]. Furthermore, the keratin proteins were neurospatially and pathologically involved in ALS. Although differential expression is often an attribute of keratin contaminants, pathological and neurospatial relevance are not.

## 5 Conclusion

In conclusion, this research made the State-of-the-Art method for diagnosing ALS with applications to other neurodegenerative diseases such as Alzheimer’s given its high accuracy on the Alzheimer’s transcriptomic dataset. This research also made the first ever correlation of KRT6A, KRT6B and KRT4 with ALS as specific biomarkers that have the potential to improve the efficacy of interventional therapies through early diagnosis and early monitoring. This research also suggests that these proteins contribute to hormonal dysregulation in ALS. Lastly, this research made the first ever correlation of PT150 with ALS as a BBB-permeable, multitarget, and neuroprotective therapeutic with abilities to assist in repairing motor neurons and regulating hormones.

## Data Availability

The whole blood ALS transcriptomic dataset is publicly available at Gene Expression Omnibus (GEO): Accession GSE112681, ID 200112681. The cerebrospinal fluid proteomics dataset is available upon reasonable request to Adam N. Trautwig of the Department of Neurology, Emory University.

## Author Information

### 5.3 Contributions

**S.D**.: Conceptualization, Methodology, Software, Validation, Investigation, Data Curation, Visualization. **A.S**.: Conceptualization, Methodology, Validation, Formal analysis, Resources, Data Curation, Writing, Visualization, Supervision. **A.D**.: Conceptualization, Validation, Investigation, Visualization. **P.K**.: Supervision.

### 5.4 Corresponding Author

Correspondence to Arnav Sharma

## References

[1] Yu, W., Yu, W., Yang, Y., & Lü, Y. (2021). Exploring the Key Genes and Identification of Potential Diagnosis Biomarkers in Alzheimer’s Disease Using Bioinformatics Analysis. Frontiers in Aging Neuro-science. https://doi.org/10.3389/fnagi.2021.602781

[2] Nguyen, H.P., Van Broeckhoven, C., & van der Zee, J. (2018). ALS Genes in the Genomic Era and Their Implications for FTD. Trends in Genetics. https://doi.org/10.1016/j.tig.2018.03.001

[3] Leblond, C.S., Kaneb, H.M., Dion, P.A., & Rouleau, G.A. (2014). Dissection of Genetic Factors Associated with Amyotrophic Lateral Sclerosis. Experimental Neurology. https://doi.org/10.1016/j.expneurol.2014.04.013

[4] Kaur, S.J., McKeown, S.R., & Rashid, S. (2016). Mutant SOD1 Mediated Pathogenesis of Amyotrophic Lateral Sclerosis. Gene. https://doi.org/10.1016/j.gene.2015.11.049

[5] The ALS Association. (2025). Genetics of ALS. Accessed March 28, 2025, from https://www.als.org/research/als-research-topics/genetics.

[6] Neumann, M., Rademakers, R., Roeber, S., Baker, M., Kretzschmar, H.A., & Mackenzie, I.R. (2009). A new subtype of frontotemporal lobar degeneration with FUS pathology. Brain. https://doi.org/10.1093/brain/awp214

[7] Nag, S., & Schneider, J.A. (2023). Limbic-predominant age-related TDP43 encephalopathy (LATE) neuropathological change in neurodegenerative diseases. Nature Reviews Neurology. https://doi.org/10.1038/s41582-023-00846-7

[8] Josephs, K.A., Murray, M.E., Whitwell, J.L., Parisi, J.E., Petrucelli, L., Jack, C.R., & Dickson, D.W. (2016). Updated TDP-43 in Alzheimer’s Disease Staging Scheme. Acta Neuropathologica. https://doi.org/10.1007/s00401-016-1537-1

[9] Rosen, D.R., Siddique, T., Patterson, D., et al. (1993). Mutations in Cu/Zn superoxide dismutase gene are associated with familial amyotrophic lateral sclerosis. Nature. https://doi.org/10.1038/362059a0

[10] Kwiatkowski, T.J. Jr., Bosco, D.A., Leclerc, A.L., et al. (2009). Mutations in the FUS/TLS gene on chromosome 16 cause familial amyotrophic lateral sclerosis. Science. https://doi.org/10.1126/science.1166066

[11] Gregory, J.M., Fagegaltier, D., Phatnani, H., & Harms, M.B. (2020). Genetics of Amyotrophic Lateral Sclerosis. Current Genetic Medicine Reports. https://doi.org/10.1007/s40142-020-00194-8

[12] Kokhlikyan, N., Miglani, V., Martin, M., Wang, E., Alsallakh, B., Reynolds, J., Melnikov, A., Kliushkina, N., Araya, C., Yan, S., & Reblitz-Richardson, O. (2020). Captum: A unified and generic model interpretability library for PyTorch. arXiv preprint. https://doi.org/10.48550/arXiv.2009.07896

[13] Hamilton, W.L., Ying, R., & Leskovec, J. (2017). Inductive representation learning on large graphs. Advances in Neural Information Processing Systems. https://doi.org/10.48550/arXiv.1706.02216

[14] Ashburner, M., Ball, C.A., Blake, J.A., Botstein, D., Butler, H., Cherry, J.M., Davis, A.P., Dolinski, K., Dwight, S.S., Eppig, J.T., Harris, M.A., Hill, D.P., Issel-Tarver, L., Kasarskis, A., Lewis, S., Matese, J.C., Richardson, J.E., Ringwald, M., Rubin, G.M., & Sherlock, G. (2000). Gene ontology: Tool for the unification of biology. Nature Genetics. https://doi.org/10.1038/75556

[15] Renton, A.E., Majounie, E., Waite, A., Simon-Sanchez, J., Rollinson, S., Gibbs, J.R., Schymick, J.C., Laaksovirta, H., van Swieten, J.C., Myllykangas, L., et al. (2011). A hexanucleotide repeat expansion in C9orf72 is the cause of chromosome 9p21-linked ALS-FTD. Neuron. https://doi.org/10.1016/j.neuron.2011.09.010

[16] DeJesus-Hernandez, M., Mackenzie, I.R., Boeve, B.F., Boxer, A.L., Baker, M., Rutherford, N.J., et al. (2011). Expanded GGGGCC hexanucleotide repeat in noncoding region of C9orf72 causes chromosome 9p-linked FTD and ALS. Neuron. https://doi.org/10.1016/j.neuron.2011.09.011

[17] Majounie, E., Renton, A.E., Mok, K., Dopper, E.G., Waite, A., Rollinson, S., et al. (2012). Frequency of the C9orf72 hexanucleotide repeat expansion in patients with amyotrophic lateral sclerosis and frontotemporal dementia: a cross-sectional study. The Lancet Neurology. https://doi.org/10.1016/S1474-4422(12)70043-1

[18] Neumann, M., Sampathu, D.M., Kwong, L.K., Truax, A.C., Micsenyi, M.C., Chou, T.T., Bruce, J., Schuck, T., Grossman, M., Clark, C.M., et al. (2006). Ubiquitinated TDP-43 in frontotemporal lobar degeneration and amyotrophic lateral sclerosis. Science. https://doi.org/10.1126/science.1134108

[19] Van Rheenen, W., Diekstra, F.P., Harschnitz, O., Westeneng, H.J., van Eijk, K.R., Saris, C.G.J., Groen, E.J.N., van Es, M.A., Blauw, H.M., van Vught, P.W.J., Veldink, J.H., & van den Berg, L.H. (2018). Whole blood transcriptome analysis in amyotrophic lateral sclerosis: A biomarker study. PLOS One. https://doi.org/10.1371/journal.pone.0198874

[20] Trautwig, A.N., Fox, E.J., Dammer, E.B., Shantaraman, A., Ping, L., Duong, D.M., Levey, A., Lah, J.J., Fournier, C.N., McEachin, Z.T., Glass, J.D., & Seyfried, N.T. (2024). Network analysis of the cerebrospinal fluid proteome reveals shared and unique differences between sporadic and familial forms of amyotrophic lateral sclerosis. bioRxiv. https://doi.org/10.1101/2024.02.29.582840

[21] Benatar, M., Zhang, L., Wang, L., Granit, V., Statland, J., Barohn, R., et al. (2020). Validation of serum neurofilaments as prognostic and potential pharmacodynamic biomarkers for ALS. Neurology. https://doi.org/10.1212/WNL.0000000000009559

[22] Target ALS. (2022). False Positives and False Negatives: How ALS Can Be Mis-diagnosed. Accessed March 16, 2025, from https://www.targetals.org/2022/02/02/false-positives-and-false-negatives-how-als-can-be-misdiagnosed/.

[23] Hawrylycz, M., Lein, E., Guillozet-Bongaarts, A., et al. (2012). An anatomically comprehensive atlas of the adult human brain transcriptome. Nature. https://doi.org/10.1038/nature11405

[24] The UniProt Consortium. (2024). UniProt: the Universal Protein Knowledgebase in 2025. Nucleic Acids Research. https://doi.org/10.1093/nar/gkae1010

[25] Thul, P.J., & Lindskog, C. (2017). The human protein atlas: A spatial map of the human proteome. Protein Science. https://doi.org/10.1002/pro.3307

[26] Tafuri, B., Milella, G., Filardi, M., Giugno, A., Zoccolella, S., Tamburrino, L., Gnoni, V., Urso, D., De Blasi, R., Nigro, S.S., & Logroscino, G. (2024). Machine learning-based radiomics for Amyotrophic Lateral Sclerosis diagnosis. Expert Systems With Appli-cations. https://doi.org/10.1016/j.eswa.2023.122585

[27] Lenail, B. (2019). NN-SVG: Publication-Ready Neural Network Architecture Schematics. Journal of Open Source Software. https://doi.org/10.21105/joss.00747

[28] Rose, T., Monti, N., Anand, N., & Shen, T. (2023). PLAPT: Protein-Ligand Binding Affinity Prediction Using Pretrained Transformers. bioRxiv. https://doi.org/10.1101/2024.02.08.575577

[29] Bugnon, M., Röhrig, U.F., Goullieux, M., Perez, M.A.S., Daina, A., Michielin, O., & Zoete, V. (2024). SwissDock 2024: major enhancements for small-molecule docking with Attracting Cavities and AutoDock Vina. Nucleic Acids Research, 52 (W1), W324–W332. https://doi.org/10.1093/nar/gkae300

[30] BioRender. (2025). BioRender - Science Illustration Tool. Accessed March 20, 2025, from https://biorender.com.

[31] Druzhilovskiy, D.S., Rudik, A.V., Filimonov, D.A., Gloriozova, T.A., Lagunin, A.A., Dmitriev, A.V., Pogodin, P.V., Dubovskaya, V.I., Ivanov, S.M., Tarasova, O.A., Bezhentsev, V.M., Murtazalieva, K.A., Semin, M.I., Maiorov, I.S., Gaur, A.S., Sas-try, G.N., & Poroikov, V.V. (2018). Computational platform Way2Drug: from the prediction of biological activity to drug repurposing. Russian Chemical Bulletin. https://doi.org/10.1007/s11172-017-1952-z

[32] Daneshafrooz, N., Bagherzadeh Cham, M., Majidi, M., & Panahi, B. (2022). Identification of potentially functional modules and diagnostic genes related to amyotrophic lateral sclerosis based on the WGCNA and LASSO algorithms. Scientific Reports. https://doi.org/10.1038/s41598-022-24306-2

[33] Morselli, L. L., Bongioanni, P., Genovesi, M., Licitra, R., Rossi, B., Murri, L., Rossi, G., Martino, E., & Gasperi, M. (2006). Growth hormone secretion is impaired in amyotrophic lateral sclerosis. Clin-ical Endocrinology, https://doi.org/10.1111/j.1365-2265.2006.02609.x

[34] Patacchioli, F. R., Monnazzi, P., Scontrini, A., Tremante, E., Caridi, I., Brunetti, E., Buttarelli, F. R., & Pontieri, F. E. (2003). Adrenal dysregulation in amyotrophic lateral sclerosis. Journal of Endocrinological Investigation, https://doi.org/10.1007/BF03349149

[35] Morice, C., Baker, D. G., Patel, M. M., Nolen, T. L., Nowak, K., Hirsch, S., Kosten, T. R., & Verrico, C. D. (2021). A randomized trial of safety and pharmacodynamic interactions between a selective glucocorticoid receptor antagonist, PT150, and ethanol in healthy volunteers. Scientific Reports, https://doi.org/10.1038/s41598-021-88609-6

